## Supplemental for "Structure of *Vibrio* phage XM1, a simple contractile DNA injection machine"

### SUPPLEMENTARY INFORMATION

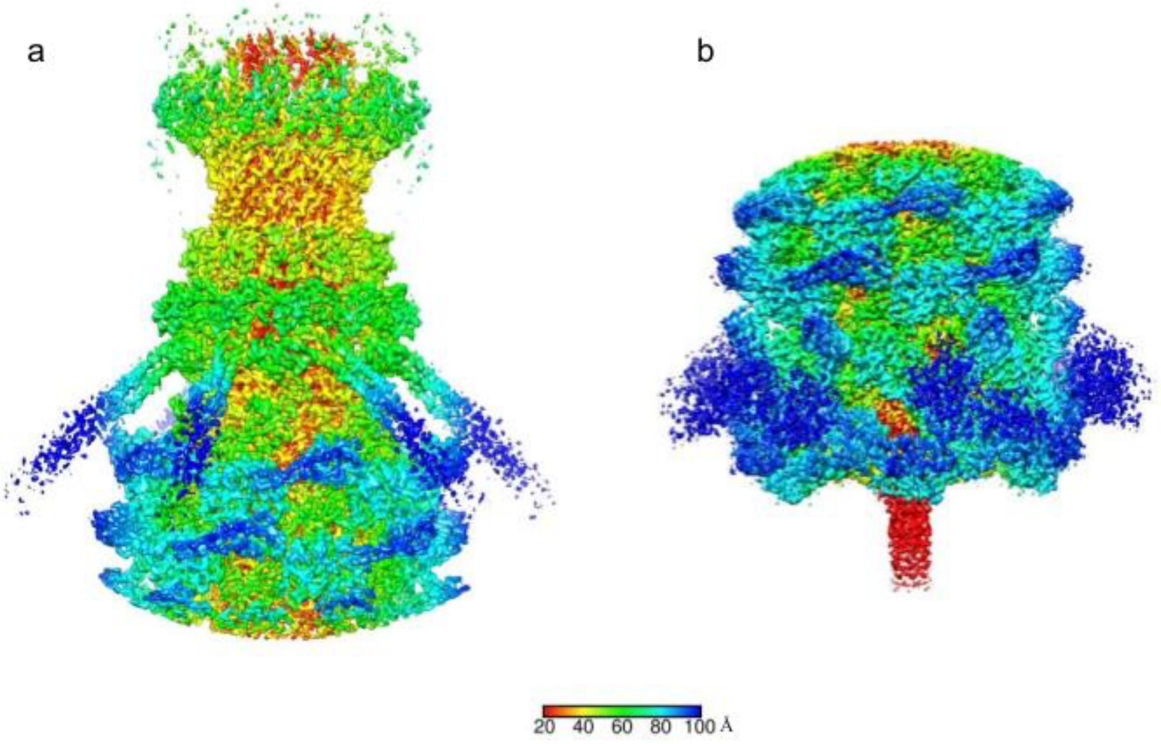


**Supplementary Fig. 1. Cryo-EM reconstructions of the XM1 neck region (a) and the baseplate region (b) calculated using 6-fold symmetry.** The density is colored according to the distance from the 6-fold axis as indicated by the color bar.

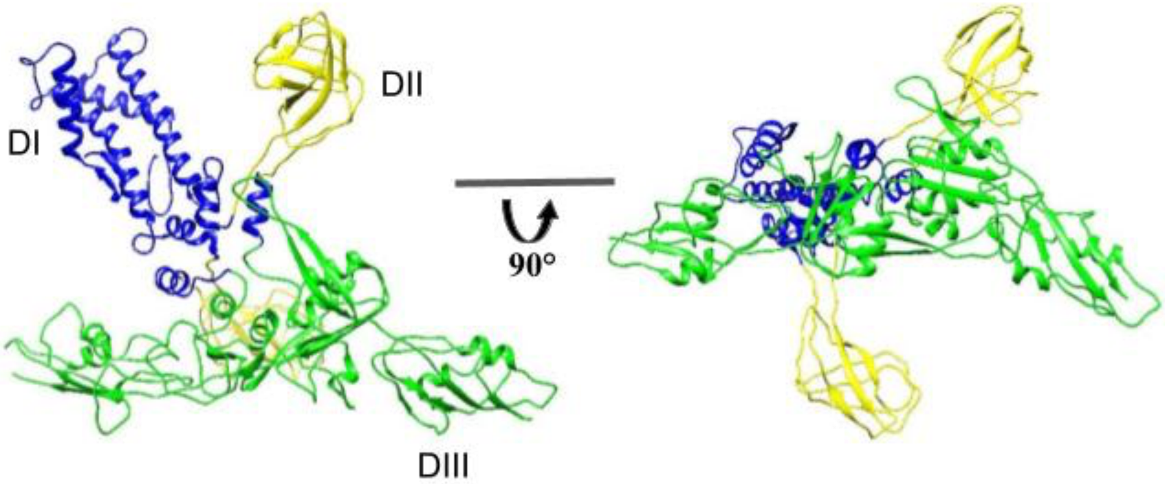


**Supplementary Fig. 2. Two views of the gp16 dimer.** Domains I, II, and III are colored blue, yellow and green, respectively. The two copies of gp16 adopt different conformations.

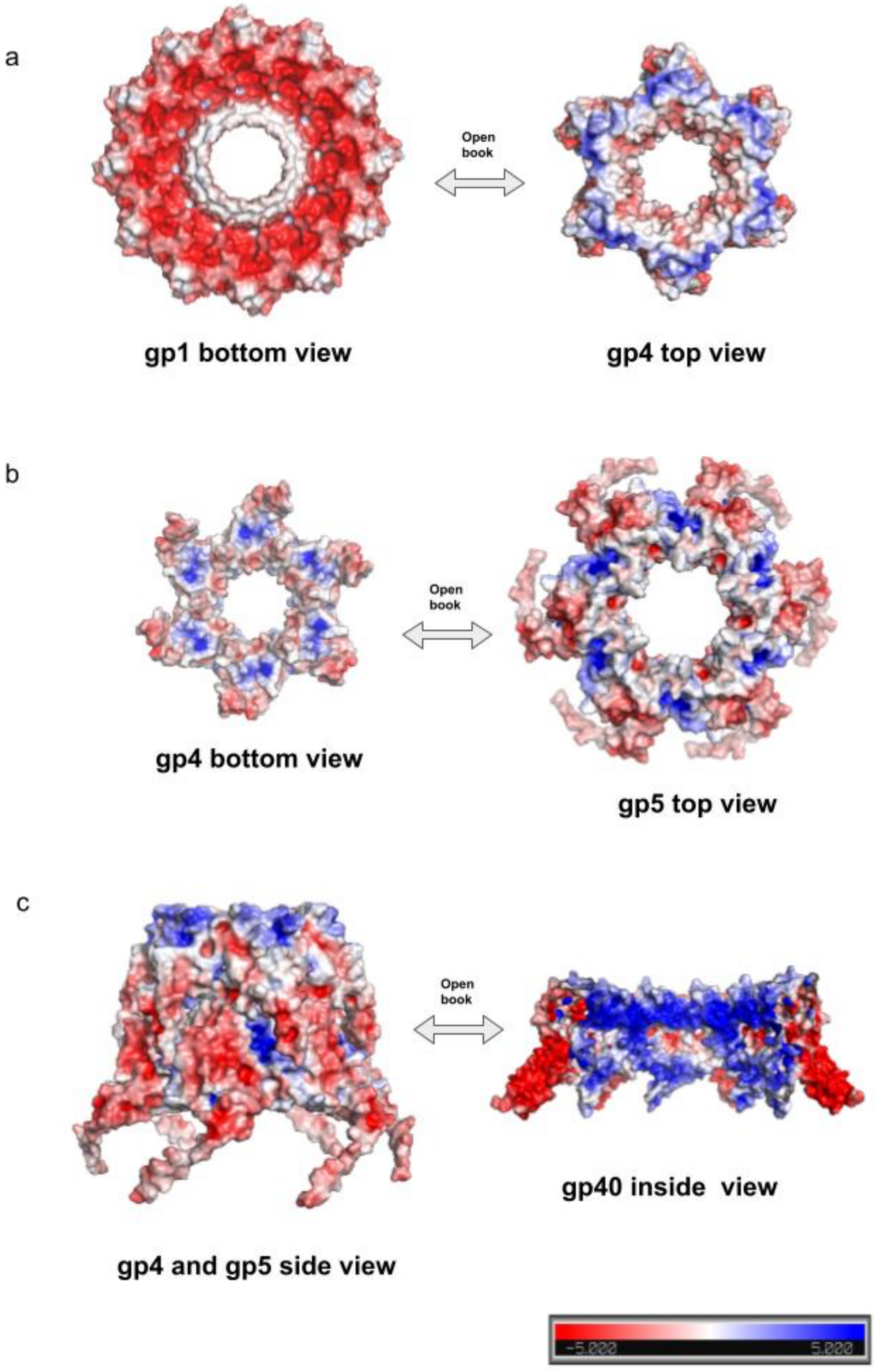
a
Open
book
gp1 bottom view
gp4 top view
b
Open
book
gp4 bottom view
gp5 top view
c
Open
book
gp40 inside view
gp4 and gp5 side view
-5.000
5.000

**Supplementary Fig. 3. Electrostatic interactions between gp1, gp4, gp5, and gp40. a.** The open book view of the molecular surfaces of gp1 and gp4 colored according to electrostatic potential. Red corresponds to a potential of -5 $kT/e^-$ and blue corresponds to a potential of +5 $kT/e^-$. This shows that the bottom surface of gp1 is negatively charged and the top surface of gp4 is positively charged. Therefore, they tightly bind to each other. **b.** The open book view of the gp4 and gp5. The surfaces are colored according to electrostatic potential. The bottom surface of gp4 and the top surface of gp5 are both positively charged and do not show strong electrostatic attraction. **c.** The side surface of the gp4-gp5 complex (shown on the left) is negatively charged. The inner surface of the gp40 ring (shown on the right) is positively charged. Therefore, assembly of the gp40 ring around the gp4-gp5 complex reinforces the interactions between gp4 and gp5. As the gp4 and gp5 proteins probably form the interface between the capsid and tail, gp40 is likely to reinforce the head-tail interactions.

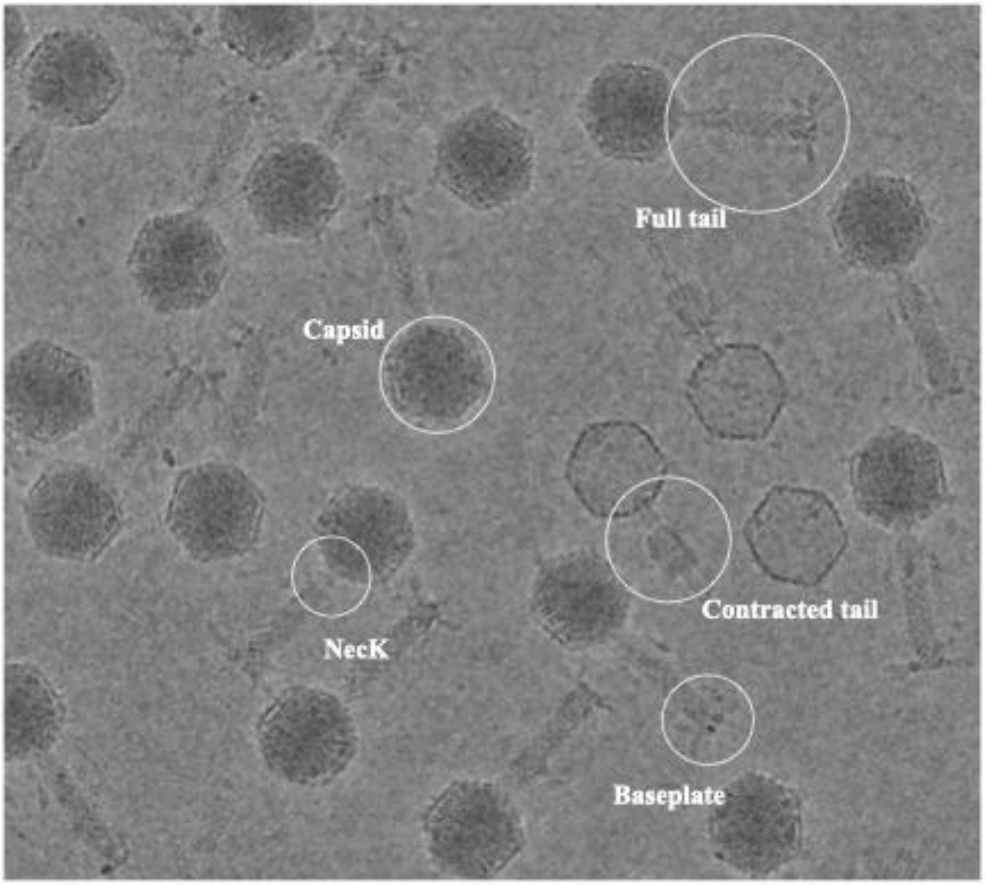


**Supplementary Fig. 4. Micrograph of the phage XM1 sample.** White circles indicate particles boxed for reconstructions.

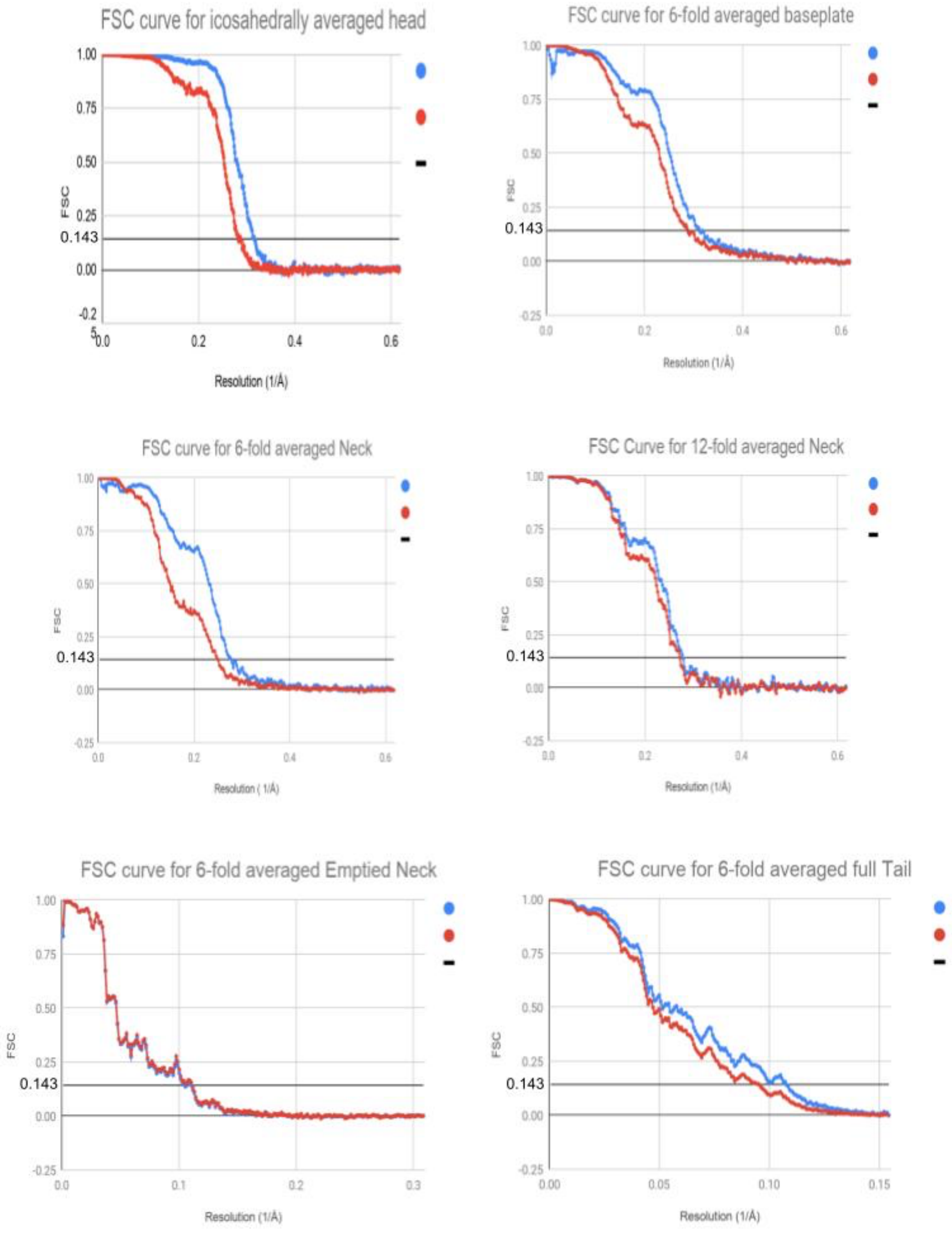


**Supplementary Fig. 5. Fourier shell correlation curves of the cryo-EM reconstructions.**

### Supplementary Table 1. Structural components of the XM1 virion.

| | # of AA built/Total AA | Predicted function | HHPRED with probability score | Dali search with RMSD | phage T4 homologue | T6SS homologue |
|---|---|---|---|---|---|---|
| gp1 | 118AA/118AA | Head-tail Connector Protein | phage Spp1 gp15 (5A21_C), 91.41 | phage Mu gp36 (5YDN), 2.9 | gp13 | - |
| gp4 | 114AA/114AA | Head-tail Connector Protein | phage Spp1 gp16 (5A21_E), 98.57 | phage Spp1 gp16 (2KCA), 3.9 | gp14 | - |
| gp5 | 160AA/161AA | Tail terminator Protein | phage Spp1 gp17 (5A21_G), 97.05 | phage Lambda gpU (3FZ2_E), 3.5 | gp15 | - |
| gp6 | 497AA/497AA | Tail Sheath Protein | R-type bacteriocin sheath protein (6GKW_A), 99.87 | R-type pyocin sheath (3J9Q_A), 3.6 | gp18 | TssB/C, VipA/B |
| gp7 | 142AA/143AA | Tail Tube Protein | R-type bacteriocin tube protein (6GKX_A), 96.73 | antifeeding prophage AFP1 (6RAP_A), 2.7 | gp19 | HCP |
| gp10 | 0/479AA | Tape Measure Protein | Phage 80a gp57 tape measure protein (6V8I_BF), 98.44 | | gp29 | - |
| gp11 | 250AA/250AA | Baseplate Organizing Protein | Pyocin R2 tail tube protein(6U5B_K), 97.13 | HCP1 (6BDC_A), 2.7 | gp48/gp53 | HCP |
| gp12 | 118AA/118AA | Baseplate Stabilizing Protein | | Rift Valley Fever Virus Envelope protein (6EGT_A), 2.8 | - | |
| gp13 | 0/376AA | Tail Hub Protein | Pyocin R2 tail hub protein (6U5H_A), 99.88 | | gp5/gp27 | VgrG |
| gp14 | 0/240AA | Tail Puncturing Protein | phageSN gp41 puncturing protein (4RU3_A), 99.89 | | gp5.4 | VgrG PAAR |
| gp15 | 110AA/117AA | Sheath initiator | phage T4 gp25 (5IW9_B), 99.75 | phage T4 gp25 (5IW9_A), 5.0 | gp25 | TssE |
| gp16 | 404AA/404AA | Baseplate Wedge Protein | phage T4 gp7 (5HX2_D), 99.97 | phage T4 gp6 (5HX2_D), 5.2 | gp6 | TssF |
| gp17 | 172AA/242AA (1-79, 107-200) | Baseplate Wedge Protein | Pyocin R2 tri1a (6U5B_w), 98.77 | phage T4 gp7 (5HX2_A), 5.2 | gp7 | TssG |

| | | | | | | |
|---|---|---|---|---|---|---|
| gp18 | 0/653AA | Tail Spike Protein | phage vb_AbaP_AS12 gp42 tailspike (6EU4_B), 95.52 | | - | - |
| gp40 | 126AA/839AA (1-126) | Collar Spike Protein | phage Det7 gp208 tailspike (6F7D_A), 99.71 | | - | - |
| gp49 | 365AA/412AA (9-337, 346-377) | Portal Protein | phage G20C portal protein (5NGD_D), 99.64 | phage G20C portal protein (4ZJN_A), 4.0 | gp20 | - |
| gp53 | 160AA/160AA | Minor Capsid Protein | | phage TW1 minor capsid protein (5WK1_K), 3.2 | - | - |
| gp54 | 294AA/324AA (31-324) | Major Capsid Protein | phage TW1 major capsid protein (5WK1_C), 100 | phage TW1 major capsid protein (5WK1_A), 3.8 | gp23/gp24 | - |

### Supplementary Table 2. Structure refinement statistics.

| | EMDB map: 22931<br><br>PDB: 7KMX | EMDB map: 22917<br><br>PDB: 7KLN | EMDB map: 22896<br><br>PDB: 7KJK | EMDB map: 22873<br><br>PDB: 7KH1 | EMDB map: 22960<br><br>-- | EMDB map: 22959<br><br>-- |
|---|---|---|---|---|---|---|
| **Data collection and processing** | | | | | | |
| Voltage (kV) | 300 | 300 | 300 | 300 | 300 | 300 |
| Electron exposure ($e^-/Å^2$) | 30 | 30 | 30 | 30 | 30 | 30 |
| Defocus range (um) | 0.5~2.5 | 0.5~2.5 | 0.5~2.5 | 0.5~2.5 | 0.5~2.5 | 0.5~2.5 |
| Pixel size (Å) | 0.82 | 0.82 | 0.82 | 0.82 | 3.24 | 3.24 |
| Symmetry imposed | icosahedral | C12 | C6 | C6 | C6 | C6 |
| Final particle images (no.) | 19625 | 10615 | 10615 | 16068 | 1104 | 6844 |
| Map resolution (Å) @FSC=0.143 | 3.2 | 3.6 | 3.6 | 3.2 | 9.1 | 10.6 |
| **Refinement** | | | | | | |
| Model composition | | | | | -- | -- |
| Non-hydrogen atoms | 24045 | 45564 | 59088 | 97344 | -- | -- |
| Protein residues | 3178 | 5748 | 7740 | 12588 | -- | -- |
| Ligands | 0 | 0 | 0 | 0 | -- | -- |
| R.M.S. deviations | | | | | | |
| Bond lengths (Å) | 0.006 | 0.007 | 0.009 | 0.009 | -- | -- |

| | | | | | | |
|---|---|---|---|---|---|---|
| Bond angles (°) | 0.634 | 0.851 | 0.933 | 0.87 | -- | -- |
| Validation | | | | | | |
| MolProbity score | 1.92 | 2.48 | 2.61 | 2.42 | -- | -- |
| Clashscore | 11.93 | 25.61 | 27.03 | 20.43 | -- | -- |
| Poor rotamers (%) | 0.12 | 0.48 | 1.45 | 0.35 | -- | -- |
| Ramachandran | | | | | | |
| Favored (%) | 95.27 | 88.71 | 89.26 | 87.1 | -- | -- |
| Allowed (%) | 4.73 | 11.29 | 10.74 | 12.71 | -- | -- |
| Disallowed (%) | 0 | 0 | 0 | 0.19 | -- | -- |
